# Dissecting Reactive Astrocyte Responses: Lineage Tracing and Morphology-based Clustering

**DOI:** 10.1101/2024.04.01.587565

**Authors:** Lina M. Delgado-García, Ana C. Ojalvo-Sanz, Thabatta K. E. Nakamura, Eduardo Martín-López, Marimelia Porcionatto, Laura Lopez-Mascaraque

## Abstract

Graphical Abstract

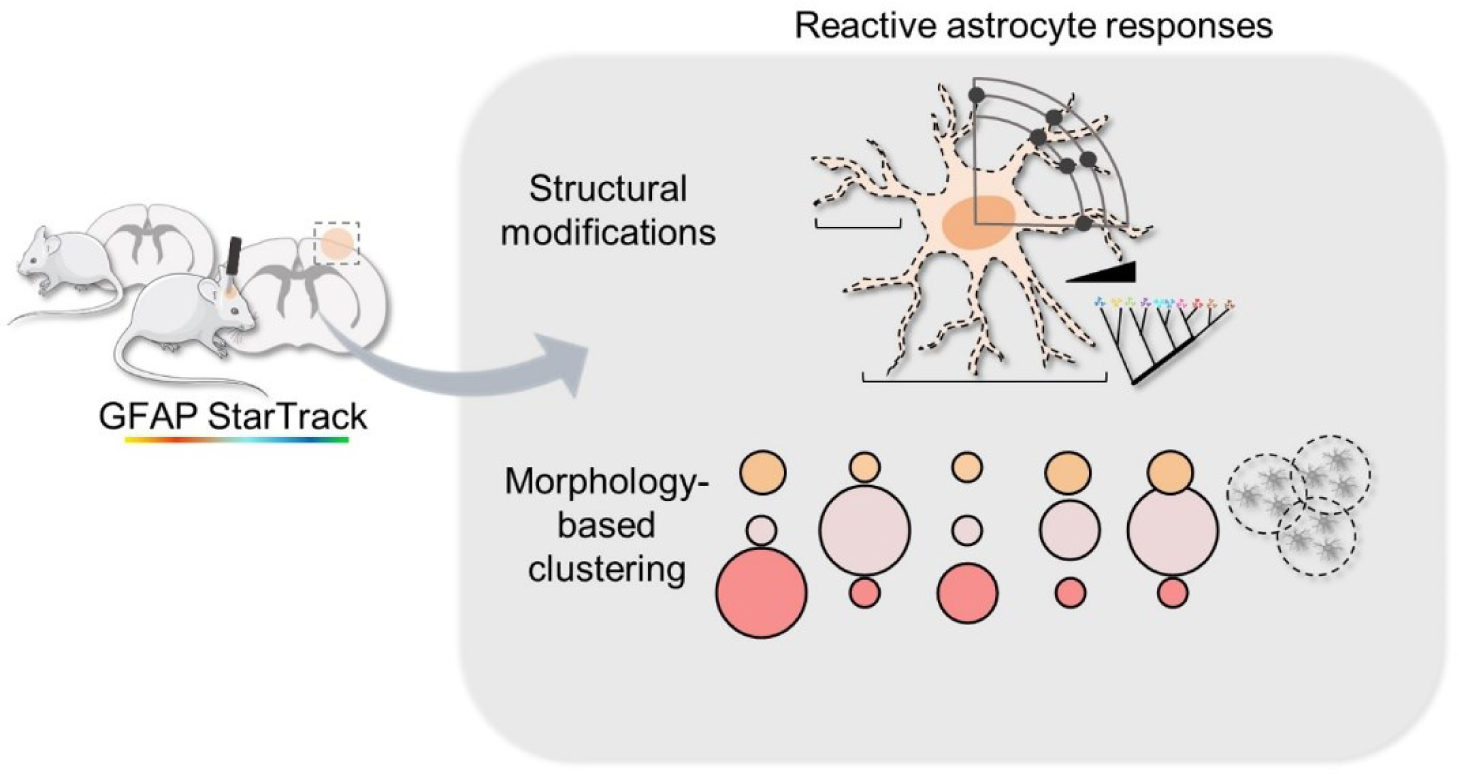

Brain damage triggers diverse cellular and molecular events, with astrocytes playing a crucial role in activating local neuroprotective and reparative signaling within damaged neuronal circuits. Here, we investigated reactive astrocytes using a multidimensional approach to categorize their responses into different subtypes based on morphology using the StarTrack lineage tracer, single-cell imaging reconstruction and multivariate data analysis. Our findings revealed three profiles of reactive astrocyte responses affecting cell size- and shape-related morphological parameters: “moderate,” “strong,” and “very strong”. We also explored the heterogeneity in astrocyte reactivity, with a particular emphasis in the spatial and clonal distribution. Our research highlights the importance of the relationships between the different astrocyte subpopulations with their reactive responses, showing an enrichment of protoplasmic and fibrous astrocytes within the “strong” and “very strong” subtypes. Overall, our study contributes to a better understanding of astrocyte heterogeneity in response to an injury. By elucidating the diverse reactive responses among astrocyte subpopulations, we pave the way for future research aimed at uncovering novel therapeutic targets for mitigating the effects of brain damage and promoting neural repair.

## Introduction

Brain damage triggers a complex cascade of cellular and molecular events aimed at restoring homeostasis and neural signaling. Among key cellular players, astrocytes develop neuroprotective or detrimental responses depending on the balance between the gain and loss of their homeostatic functions (Escartin et al., 2019; Verkhratsky et al., 2023). At the molecular lever, astrocyte reactivity results in the reorganization of their intermediate filaments, a cytoskeletal network formed by relevant glial proteins such as GFAP, vimentin, and nestin. Increased GFAP content is responsible for characteristic astrocyte responses such as resistance to mechanical stress, formation of glial scars, vesicle trafficking, and autophagy (Eliasson et al. 1999; Wang et al. 2018; Escartin et al., 2019; Potokar et al. 2020).

However, major questions regarding astrocyte morphology changes in relation to injury such as how astrocytes acquire their shapes, whether and how these changes alter neuron-glia interactions, or whether astrocyte changes contribute to the disease causation and progression, remain unknown (Baldwin et al., 2023). Furthermore, the morphological modifications among astrocyte subpopulations, known generally as “astrocyte reactivity”, into the local neuroprotective and reparative signaling within damaged neuronal circuits, continue to be largely unstudied. By elucidating the role of structural modifications in reactive astrocytes, we can advance our knowledge on the biology of glial cells and contribute to the exploration of brain reparative mechanisms at the single-cell level.

The StarTrack lineage tracer (García-Marqués and López-Mascaraque, 2013) is a powerful method for genetically and permanently labeling astrocyte progenitors and their progenies, enabling the identification of mature cortical astrocyte clones (reviewed in Bribián et al., 2015 and Figueres-Oñate et al., 2021). This method is also an effective tool to study morphological and biochemical changes of cortical astrocytes after brain damages such as experimental autoimmune encephalomyelitis (EAE) and traumatic brain injury (TBI; Bribián et al., 2018; Martín-López et al., 2013; Barriola et al., 2020; Götz et al., 2020). Previously, we observed that most cortical astrocyte clones exhibited a strong reactive phenotype in response to injury, indicating the presence of intrinsically heterogeneous reactive morphological responses (Martín-López et al., 2013). In this work we took a step further and used the StarTrack method, in conjunction with single-cell imaging reconstruction and multivariate data analysis, to elucidate the underlying reactive responses among different subpopulations of cortical astrocytes. We categorized these astrocyte responses using morphological parameters in a TBI model and provide a comprehensive frame to classify reactive astrocytes based on the intensity of their response to the lesion. Our research involved a comprehensive examination of single-cell modifications in key astrocytic components, coupled with the development of a structure-based classification system for reactive astrocyte responses.

## Results

### Morphological characterization of cortical reactive astrocytes in response to Traumatic Brain Injury (TBI)

In this study, we employed the StarTrack genetic lineage tracing method to investigate the heterogeneous reactive responses of cortical astrocytes in a model of traumatic brain injury (TBI). The StarTrack method allows the stochastic expression of 6 different fluorophores expressed in each of the two main cellular compartments, the cytoplasm, and the nucleus, giving rise to a total of 12 possible combinatorial expressions of colors that provide a unique ID color code to identify astrocyte clones (García-Marqués et López-Mascaraque, 2013). Here we analyzed structural modifications of control and reactive astrocytes subpopulations and established morphology-based clusters to categorize astrocytes reactive responses (Fig.1A).

**Figure 1.**
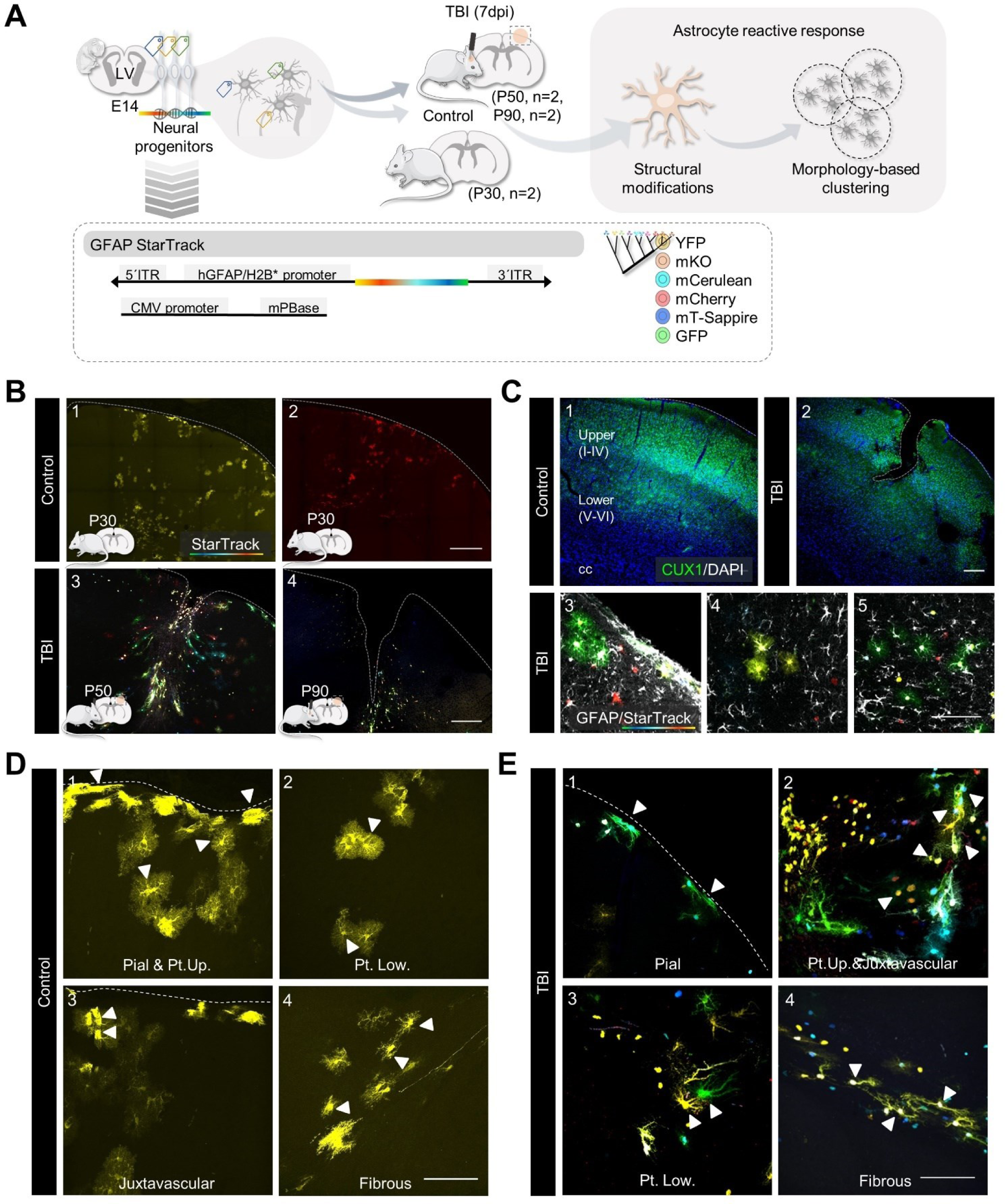
StarTrack reactive astrocyte subpopulations in a model of traumatic brain injury (TBI). A. Experimental design. Mice at embryonic day 14 (E14) were *in utero* electroporated with StarTrack. At postnatal day 50 (P50, n=2) or 90 (P90, n=2) mice were submitted to a model of TBI at the somatosensory cortex. Two control mice at postnatal day 30 were included (P30, n=2). Seven days post-injury (7dpi) we analyzed structural modifications and established a morphology-based clustering method to categorize the reactive responses of astrocytes. B. Representative images of the somatosensorial cortex of control (1,2) and TBI (3,4) StarTrack mice. C. Representative images of CUX1/DAPI immunohistochemistry. CUX1 was used to delineate upper cortical layers (II-IV; 1,2). Representative images of StarTrack astrocytes, and GFAP immunohistochemistry (3,4,5). The images are from TBI mice. D. Astrocyte subpopulations along the corpus callosum (cc), cortex (Cx) and *pia mater* (PM) from control (D) and TBI (E) mice. We identified pial (1), protoplasmic upper and lower layers (1,2,3), juxtavascular (3) and fibrous (4). Abbreviations: TBI = Traumatic brain injury; Pt. Up. = Protoplasmic upper layers; Pt. Low. = Protoplasmic lower layers. Scale bar B, left, 200 μm and right; C, 1,2, 200 μm and 4,5,6, 100 μm; D, E, 100 μm.

**Figure 2.**
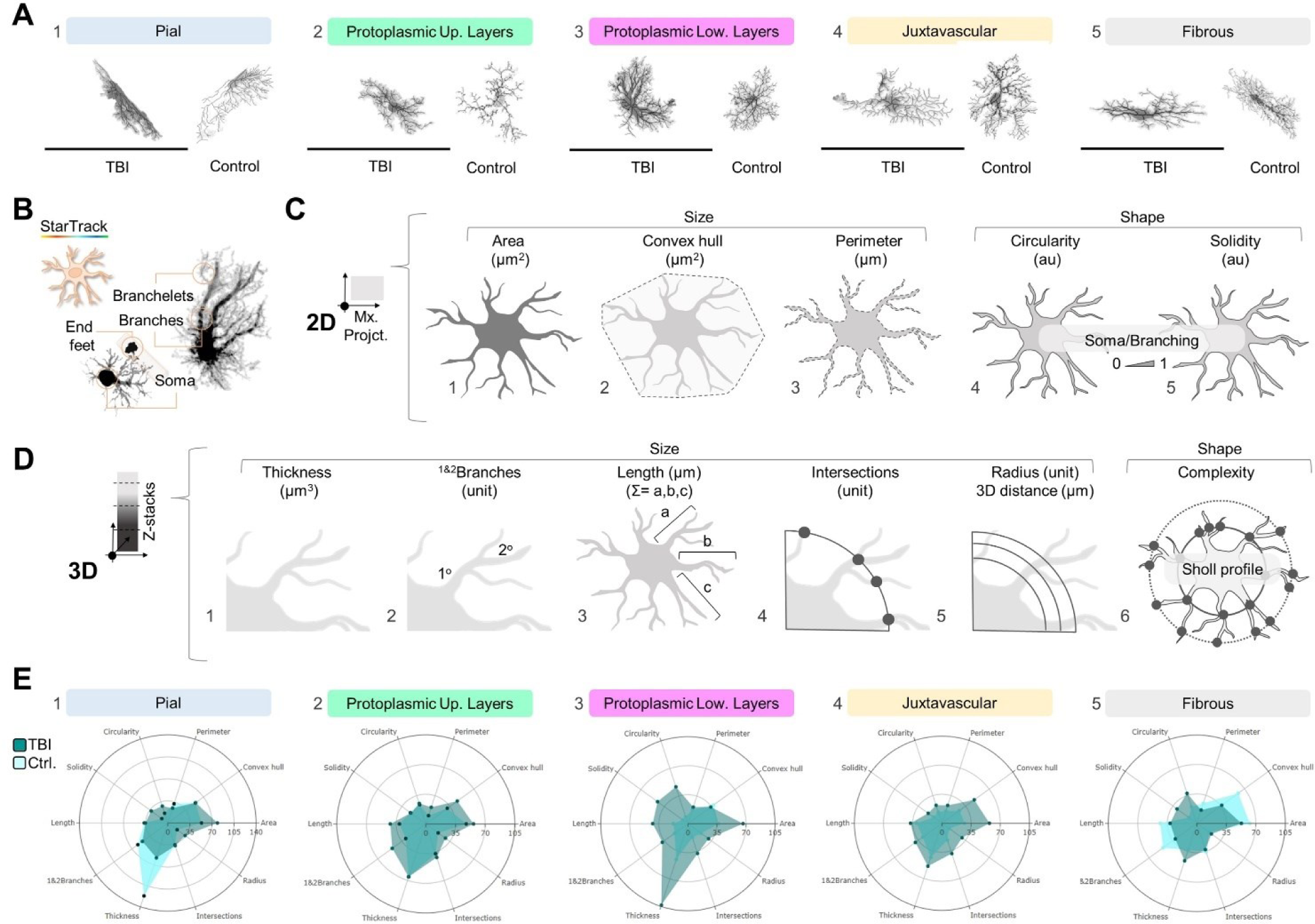
Two- and three-dimensional (2D and 3D) morphometric analysis of astrocyte subpopulations. A. Representative images of control and reactive astrocytes subpopulations: pial (1; TBI, n = 15 cells; control, n = 10 cells); protoplasmic upper layers (2; TBI, close n = 19 cells, far n = 15 cells; control, n = 10 cells); protoplasmic lower layers (3; TBI, close n = 13 cells, far n = 22 cells; control, n = 10 cells); juxtavascular (4; TBI, n = 19 cells; control, n = 10 cells); and fibrous (5; TBI, n = 15 cells; control, n = 10 cells). Sample size (n) across astrocyte subpopulations. B. Representative images of single-cell major astrocytic components-soma, primary and secondary branches, and end feet-across astrocyte subpopulations. C. Graphic representation of 2D size- and shape-related parameters: area (1; µm^2^), convex hull area (2; µm^2^), and perimeter (3; µm); 2D-shape related parameters: circularity (4; 4π(“area”)⁄(“perimeter”)^2^) and solidity (5; “area”⁄ “convex hull area”). D. Graphic representation of 3D-size and shape-related parameters: thickness (1; relative to 0.05 threshold, µm^3^), 1&2 branches (2; unit), length (3; µm), intersections (4; unit), and radius or 3D distance (5; unit, µm); and 3D-shape related parameter, complexity (6; “intersections” as a function of “3D distance”). E. CV radar graphs across astrocyte subpopulations: pial (1), protoplasmic upper layers (2), protoplasmic lower layers (3), juxtavascular (4), and fibrous (5). Astrocytes subpopulations and parameters were identified as equally significant sources of variability. Abbreviations: TBI = Traumatic brain injury; Pt.Up. = Protoplasmic upper layers; and Pt.Low. = Protoplasmic lower layers. Scale bar 100 μm.

First, we identified distinct astrocyte subpopulations in both control (Fig. 1B, 1-2) and TBI (Fig. 1B, 3-4) groups across the corpus callosum (cc), cortex (Cx) and *pia mater* (PM). To differentiate these subpopulations through the thickness of Cx, we used CUX1 immunolabeling to distinguish between upper (layers II-IV) and lower (layers V-VI) cortical astrocytes (Fig.1C, 1-2). Additionally, we delineated their location within the injury site using GFAP immunolabeling, which separated the reactive gliosis area into the “Area 1: contusion core”, and the surrounding normal-appearing tissue termed “Area 2: pericontusional” (Fig.1C, 3-5 and Fig.3A). Subsequently, we classified StarTrack labelled astrocytes from both control (Fig 1D) and TBI (Fig.1E) groups, based on their localization and morphological properties of their soma and primary branches as follows: 1) pial astrocytes (Fig.1D, 1, and Fig.1E, 1, arrowheads), also known as marginal or perimeningeal astrocytes, located at the surface and in direct contact with the PM; 2) protoplasmic astrocytes (Fig. 1D, 1-2, and Fig.1E, 2-3, arrowheads), distributed across layers I to VI; 3) juxtavascular astrocytes (Fig. 1D, 3, and Fig.1E, 2, arrowheads), attached to cortical blood vessels; and 4) fibrous astrocytes in the *cc* (Fig. 1B, 4, and Fig.1E, 4, arrowheads).

**Figure 3.**
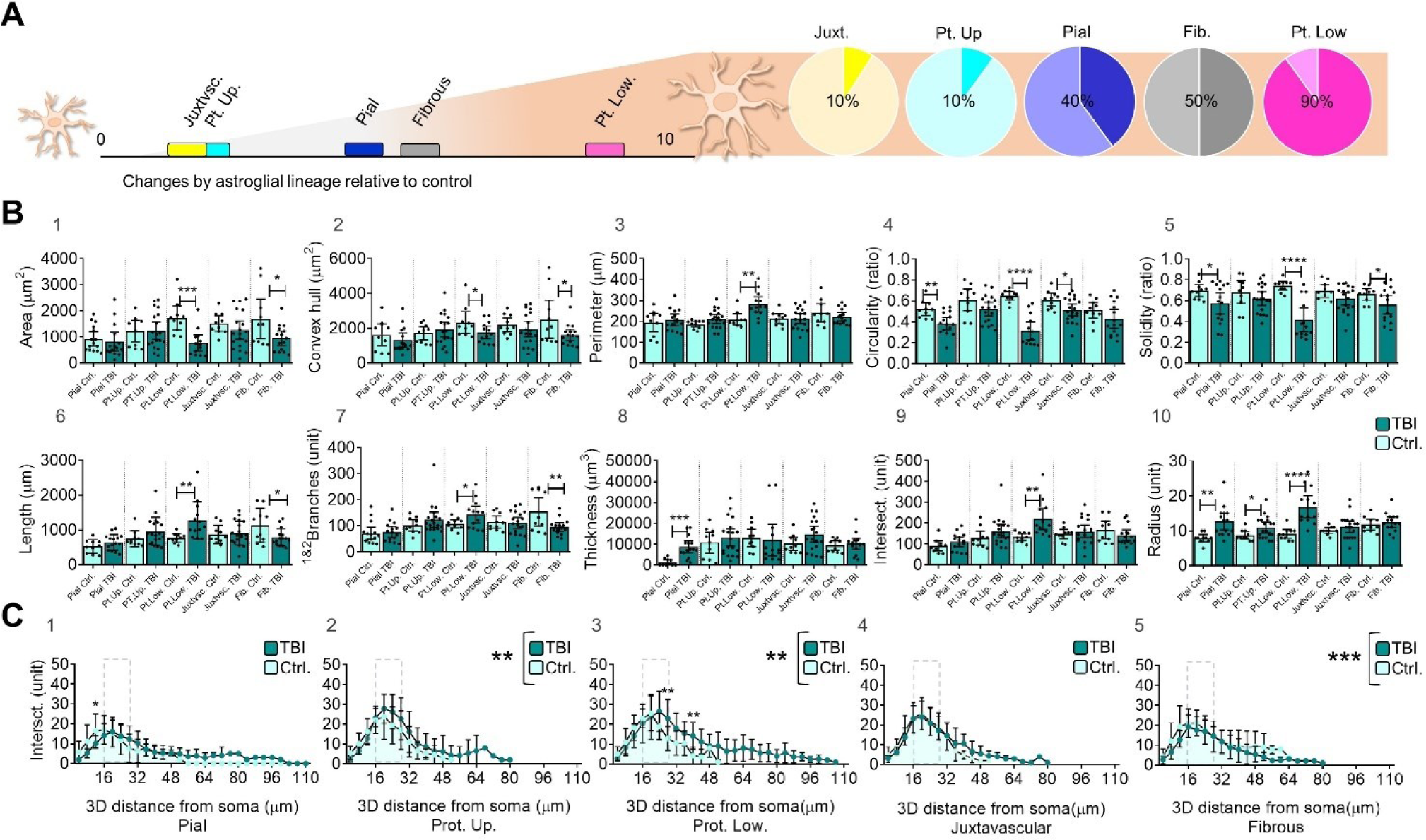
Comparison between control and reactive (TBI) astrocytes. A. Proportion of parameters with significant changes across astroglial subpopulations. Lower layer protoplasmic astrocytes exhibited a significant reactive response in nine out of the ten parameters evaluated. B. Graph bar of 2D and 3D size- and shape-related parameters in control and TBI reactive astrocytes: area (1), convex hull area (2), perimeter (3), circularity (4), solidity (5), length (6), 1&2branches (7), thickness (8), intersections (9), radius (10). The graph shows the samplés distribution, mean and 95% CI. C. XY graph of complexity (Sholl intersections profile) shows the number of intersections along the 3D soma distance. Sholl profile include three segments: an *initial growth segment* extending approximately 12 to 24 μm from the soma, mainly composed by primary branches; followed by a *peak segment* ranging between 16 to 28 μm from the soma (dashed lines) that include the highest number of intersections and their *critical-distance from the soma-value*; and a *final decline segment*, between 24 to 110 μm to the edge of the last branch. The graph shows the mean and standard deviation. Datasets on graphs in absolute values; pial, protoplasmic upper layers, protoplasmic lower layers, juxtavascular and fibrous, control n = 10 cells; pial TBI n = 15 cells; protoplasmic upper layers, TBI n = 19 cells; protoplasmic lower layers, TBI n = 13 cells; juxtavascular TBI n = 19 cells; fibrous TBI n = 15 cells. Comparison between control and reactive astrocytes was performed by unpaired T test, **** p ≤ 0.0001, *** p ≤ 0.001; ** p ≤ 0.01; * p ≤ 0.05. Abbreviations: TBI = Traumatic brain injury; Pt. Up. = Protoplasmic upper layers; and Pt. Low. = Protoplasmic lower layers.

**Figure 4.**
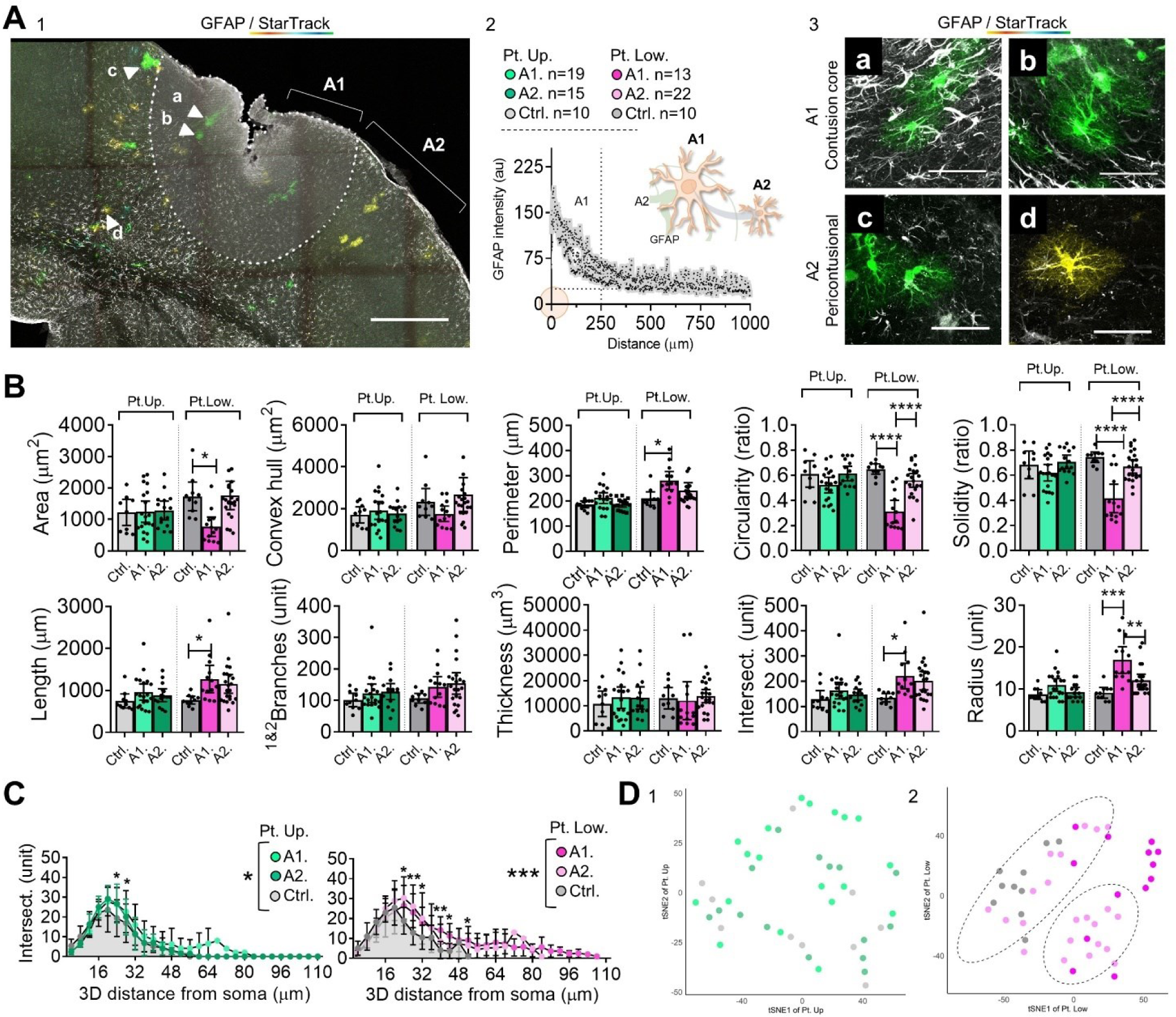
Protoplasmic reactive astrocyte distribution along TBI areas and morphometric analysis. A. Representative image of TBI main areas according to GFAP immunolabeling; Pt. Up and Pt.Low (1). Sample size and GFAP fluorescence-intensity profile (2); and representative images of GFAP/StarTrack reactive astrocytes at contusional core (A1; a, b) and pericontusional (A2; c, d) areas (3). The images are from the somatosensory cortex of TBI mice. Fluorescence-intensity profiles revealed an enriched GFAP area (A1) extending approximately 250 μm along the core, followed by a moderate GFAP-normal appearing area (A2). B. Graph bar of 2D and 3D size- and shape-related parameters in control, A1 and A2 protoplasmic (upper and lower layers) reactive astrocytes. The graph shows the sample distribution, mean and 95% confidence interval (CI). C. XY graph of complexity (Sholl intersections profile) shows the number of intersections along the 3D soma distance. The graph shows the mean and standard deviation (SD). D. T-distributed Stochastic Neighbor Embedding (T-SNE) plot and clustering analysis of upper-(1) and lower-(2) layers protoplasmic astrocytes. Datasets on graphs in absolute values; protoplasmic upper layers, control, n = 10 cells, A1, n = 19 cells, A2, n = 15 cells; protoplasmic lower layers, control, n = 10 cells, A1, n = 13 cells, A2, n = 22 cells. Comparison between control, A1 and A2 reactive astrocytes was performed by ANOVA test, **** p ≤ 0.0001, *** p ≤ 0.001; ** p ≤ 0.01; * p ≤ 0.05. Scale A1, 250 μm and A2, 50 μm. Abbreviations: Pt. Up. = Protoplasmic upper layers; and Pt. Low. = Protoplasmic lower layers.

Our analyses included morphological reconstructions of 118 StarTrack reactive astrocytes (n=4, Fig.2A, 1-5 TBI), and 45 StarTrack astrocytes that were used as controls (n=2, Fig.2A, 1-5 Control). For each cell, we identified major astrocytic components, soma, and primary and secondary branches (branchlets), to analyze their morphological profile (Fig.2B). Thus, we employed two- and three-dimensional (2D and 3D) image projections (Figs. 2C and D) to assess size- and shape-related parameters such as “cell body area” (μm^2^; Fig.2C, 1), “convex hull area” (μm^2^; Fig.2C, 2), “perimeter” (μm; Fig.2C, 3), “circularity” (4π(“area”)/(“perimeter”)^2^; Fig.2C, 4), “solidity” (“area”/”convex hull area”; Fig.2C, 5), total “thickness” (μm^3^; Fig.2D, 1), “1&2 branches” (unit; Fig.2D, 2), total branches “length” (μm; Fig.2D, 3), “intersections” (unit; Fig.2D, 4), “radius” or “3D distance” (unit, μm; Fig.2D, 5) and “complexity” (Fig.2D, 6). Altogether, this dataset let us to explore astrocyte territories and branching: parameters such as “area”, “perimeter” and “convex hull area”, measured the size of the soma and branches of astrocytes (Fig. 2C, 1-3), while parameters such as “circularity” and “solidity” (Fig. 2C, 4-5) measured their general shape variation. These shape-related parameters use ratios that compared the “area” of each astrocyte with its “perimeter”, in the case of “circularity”; or their “convex hull area”, in the case of “solidity”. Greater “perimeter” and “convex hull area” values led to a smaller “circularity” and “solidity” ratios that we interpreted as “less regular” and “less dense or spongiform” astrocyte morphologies. The parameter “thickness” (Fig 2D, 1) measured the astrocyte volume, while the parameters “1&2 branches” and “length” (Fig. 2D, 2-3) mainly determined cell branching. In addition, we also included Sholl analysis metrics such as “intersections”, “radius” or “3D distances”, and “complexity” or “Sholl intersections profile” (Fig. 2D, 4-6).

By analyzing these datasets of parameters, we constructed radar graphs with coefficient of variation values (CVs) for each parameter to gain insights into the biological trends among control and TBI groups (Fig. 2E). Interestingly, we found that both the distinct astrocyte subpopulations and morphological parameters were equally source of variability. Astrocyte variability ranged from 7.70% (CV <10%, good) in control protoplasmic solidity (lower layers, Fig. 2E, 3) to 121.8% (CV>35%, bad) in pial thickness (Fig. 2E,1). Specifically, subpopulations that exhibited acceptable interindividual variability included control protoplasmic (upper and lower layers; CV <35%, acceptable; Fig. 2E, 2-3), and juxtavascular astrocytes (Fig. 2E, 4), as well as TBI fibrous reactive astrocytes (CV <35%, acceptable; Fig. 2E, 5). Parameters with acceptable CV included “perimeter”, “circularity”, “solidity”, “length”, “intersections”, and “radius” and were efficient in detecting significant changes between control and TBI groups (Fig. 2E).

Comparisons between control and TBI groups showed that lower layer protoplasmic astrocytes exhibited significant changes in almost all size- and shape-related parameters (“area”, “convex hull”, “perimeter”, “circularity”, “solidity”, “length”, “branches”, “intersections”, and “radius”; Fig.3A; and Fig.3B, 1-10). Contrarily, upper layer protoplasmic astrocytes only changed in size (“radius”, also interpreted as the “3D distance”, Fig.3A; and Fig.3B, 1-10). Distinct responses were also observed when comparing between juxtavascular, pial and fibrous astrocytes. Thus, juxtavascular astrocytes showed only a significant change in shape (“circularity”; Fig.3A; and Fig.3B, 1-10), while pial and fibrous reactive astrocytes showed changes in both, shape- and size-related parameters (for pial: “circularity”, “solidity”, “thickness” and “radius”; for fibrous: “area”, “convex hull area”, “solidity”, “length”, and “branches”-Fig.3A and Fig.3B, 1-10). Upon conducting a more comprehensive analysis of the datasets in both the control and TBI groups, we found that under physiological conditions (control group), only the pial lineage exhibited a distinct profile among astrocyte subpopulations. However, following TBI, lower layers protoplasmic astrocytes displayed differential profiles in shape- and size-related parameters across astrocytes subpopulations (data not shown).

Further analysis of “complexity” (Sholl intersections profile) allowed the identification of three distinct segments in astrocyte branching (Fig. 3C, 1-5). These segments included: 1) an *initial growth segment* extending approximately 12 to 24 μm from the soma, mainly composed by primary branches; 2) a *peak segment* ranging between 16 to 28 μm from the soma (dashed lines) that included the highest number of intersections and their *critical-distance from the soma-value*; and 3) a *final decline segment*, with a size ranging between 24 to 110 μm to the edge of the last branch. Comparisons between control and TBI groups showed an effect in astrocyte branching density and spatial distribution for protoplasmic (upper: Fig. 3C, 2 and lower: Fig. 3C, 3) and fibrous (Fig. 3C, 5) astrocytes. In particular, protoplasmic reactive astrocytes increased their complexity while fibrous reactive astrocytes showed an opposite response (Fig.3C). Collectively, our findings revealed distinct profiles among astrocytes subpopulations, highlighting specific size- and shape-related alterations in response to the brain damage. The distinct morphological alterations, in upper- and lower-layers protoplasmic astrocytes, lead us to further investigate the relationship between astrocyte reactivity and their spatial distribution.

### Influence of the distance to the lesion in reactive astrocytes morphologies

Protoplasmic astrocytes were analyzed based on their location within the cortical layers (see above, Fig. 1C, 1-2) and distance to the injury, determined by a GFAP immunolabeling that separated the area of reactive gliosis, named as “Area 1: contusion core”, from that of normal-appearing tissue named as “Area 2: pericontusional” (Fig.1C, 3-5 and Fig.4A). Areas 1 and 2 were stablished using an image analysis extending 1000 μm from the injury site revealed GFAP fluorescence-intensity profiles that led us to define two main areas, one representing the contusion core (Area 1, A1, Fig. 4A,1-2) with increased GFAP-intensity profile (extending approximately 250 μm from the injury site) and one immediately adjacent to the contusion core, the pericontusional area (Area 2, A2) with less GFAP intensity-profile (Fig. 4A, 2). A1 was characterized as a reactive gliosis region, while A2 had a GFAP-normal-appearing profile (Fig. 1C, 3-5 and Fig.3A, 3).

Our analysis revealed that only A1 lower-layers protoplasmic astrocytes showed distinct reactive responses compared to A2 astrocytes (Fig.3B) that included morphological alterations in both size- and shape-related parameters, such as “area”, “perimeter”, “circularity”, “solidity”, “length”, “intersections”, and “radius” (Fig. 4B). Interestingly, the analysis of “complexity” (Fig. 4C) showed a group effect for both (Ctrl.-A1/Ctrl.-A2), upper- and lower-layers protoplasmic reactive astrocytes, particularly lower-layers astrocytes which displayed larger 3D distances, and differentially complexity-curve shapes with more regular decline as they moved away from the soma (Fig.4C). Pairwise analysis of each shape- and size-related parameters, and visualization of the cell population in T-distributed Stochastic Neighbor Embedding (T-SNE. Fig. 4D) plots, revealed that A1 and A2 upper layers protoplasmic control and reactive astrocytes were randomly distributed in the plot (Fig. 4D, 1) unlike most A1 lower layers protoplasmic astrocytes. On the other hand, A2 lower layers protoplasmic reactive astrocytes were distributed next to controls, indicating a similarity in their morphological profiles (Fig.4D, 2). Consistent with these findings, a further clustering analysis based on TSNE distribution (Fig. 4D, 2), resulted in grouped A2 lower layers protoplasmic reactive astrocytes with controls.

### Reactive response landscape among astrocyte lineages

Lastly, we integrated size- and shape-related parameters to construct morphology-based clusters of reactive responses and explored astrocyte lineage participation (Fig. 5). Initially, we applied a multimodal index analysis (MMI, Suppl.Fig. 1A) to guide the selection of parameters for cell clustering, focused on those with MMÍs values greater than 0.55, such as “convex hull area”, “perimeter”, “length”, “thickness”, “intersections”, and “radius” (Suppl. Fig.1A). Subsequent hierarchical clustering (HC, Fig.5A), and principal component analysis (PCA, Fig.5B) enabled the partition of the dataset into clusters labeled from A to H and their subsequent categorization as reactive responses (Fig.5A-B and Suppl. Fig.1B). Analysis of variance (ANOVA) confirmed how clusters A-B and C-D exhibited similar parameter profiles, while clusters E, F, G, H displayed greater variability (Fig.5C and Suppl. Fig.1B). Principal component 1 (PC1, 63.6%) and Principal component 2 (PC2, 17.6%) collectively explained 81.2% astrocyte variation in a bidimensional distribution, with optimal correlation values among parameters (Fig.5B and Suppl. Fig.1C, 1). PC1 included measurements of the size-related parameters “length”, “intersections”, and “perimeter”; PC2 predominantly received contributions from “thickness” and “radius”. PC biplot displayed how selected morphological parameters influenced and contributed with the direction of the datasets (Suppl. Fig.1C, 1). Upon conducting a PC biplot on relevant datasets, it became evident that the distribution of the plot is influenced by upper- and lower-layers protoplasmic astrocytes (Suppl. Fig.1C, 2).

**Figure 5.**
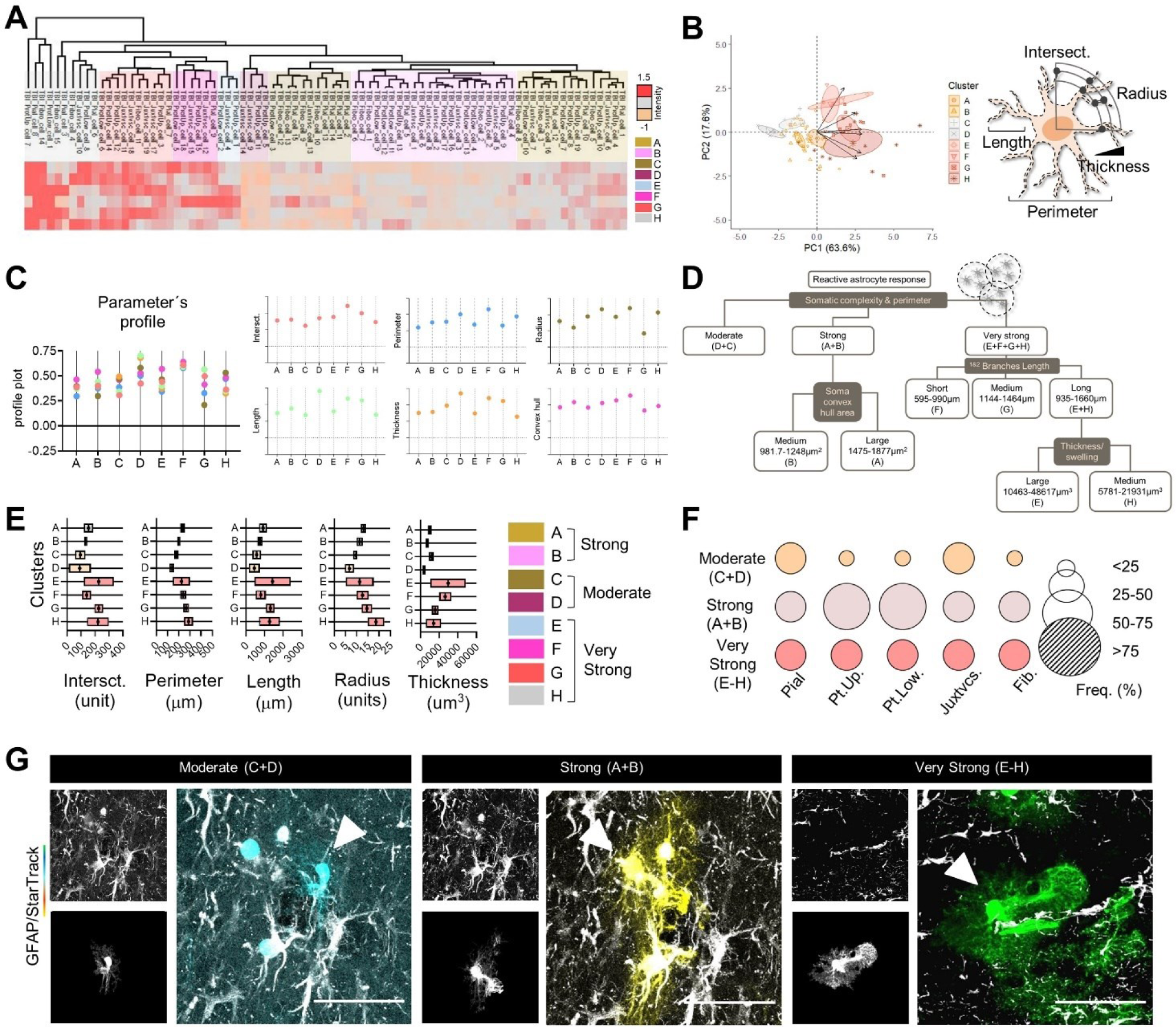
Clustering of reactive astrocytes responses. A. Heatmap showing the hierarchical clustering (HC) of reactive astrocytes subpopulations. B. Principal component analysis (PCA) plot with confidence ellipses of reactive astrocytes and clusters distribution; graphical representation of selected parameters. C. Plot profile (normalized mean) of selected parameters among clusters. D. Logical tree of reactive astrocytes response-clusters according to their somatic and branching complexity. E. Floating-bars showing the mean and 95% CI for each cluster. F. Bubble graph of astrocytes subpopulation frequency among reactive responses. G. Representative images of reactive astrocytes responses for juxtavascular subpopulations. Datasets on PCA, parameters profile and cell enrichment graphs in relative values, data sets on somatic and branching complexity graphs in absolute values. Pial n = 15 cells; protoplasmic upper layers, n = 19 cells; protoplasmic lower layers, n = 13 cells; juxtavascular n = 19 cells; fibrous n = 15 cells. Scale bar 50 μm.

Next, we built a dendrogram (logical tree) to establish diversification nodes for astrocyte reactive responses based on astrocyte clusters, individual features, and type of reactive responses (Fig.5A-D). First and second-level clustering nodes included 95% confidence interval (CI) of each parameter to suggest cut-off values (Fig.5D-E). Parameters such as “perimeter”, “intersections”, and “radius” determined the first level nodes of the tree, while parameters such as “length (branches length)” and “convex hull area (soma convex hull)” determined second- and third-level branches nodes. The parameter “thickness (swelling)” determined the fourth-level branches and terminal nodes (Fig.5D). For the first-level nodes, we classified the reactive astrocytes responses into 3 group nodes: the “moderate” node, which included astrocytes from clusters C and D (C+D); the “strong” node, including clusters A and B (A+B); and the “very strong” node that clustered the E to H astrocyte groups (E, F, G, H) (Fig.5D). Clusters A and B could be further subdivided into in “medium” and “large” second level nodes based on their soma convex hull values, while clusters E to H, (E, F, G, H) could be subdivided into “short”, “medium”, and “long” based on their processes length, and into “large” and “medium” based on their thickness (swelling) values. Interestingly, the reactive response type “very strong” consisted of clusters with greater variability (E, H), primarily characterized by extreme length and thickness values (Fig.5D).

We also examined astrocyte diversity within each reactive response (Fig. 5F). Astrocytes subpopulation frequency revealed that the “strong” type of reactive response (A+B) had the highest proportion of protoplasmic astrocytes from both upper and lower layers, while the “very strong” type exhibited similar lineage participation (Fig 5F and Suppl. Fig.3D). This diversity was clearly visible with the StarTrack method and astrocytes reconstructions, which allowed us to group the different responses based on similar morphologies (Fig.5G and Suppl. Fig.1D, 1-2). For example, the reconstruction and categorization of reactive astrocytes in two TBI mice (A147 and A117 mice; aged 50, and 90 days respectively) revealed 2 main response models: the first model (Suppl. Fig.1D, 1) experienced an enrichment of a “strong” and “very strong” type of reactive response, primarily from pial, upper-layers protoplasmic and fibrous lineages; the second model (Suppl. Fig.1D, 2) displayed proportional types of reactive responses, each enriched with pial (moderate), protoplasmic low (strong) and juxtavascular (very strong) lineages.

Finally, we conducted an exploratory analysis to characterize the morphology of the different astrocyte responses grouped in our dendrogram using one TBI mouse (A147). This analysis integrated clonal analysis, the categorization of reactive responses in astrocyte subpopulations, and their spatial distribution. Our findings demonstrated that similar reactive responses were proximal and in a similar distance to the injury (Fig.6A). In addition, a clonal analysis of 281 astrocytes corresponding to 35 clones sharing the same fluorophores combination in both, the nucleus, and the cytoplasm (Fig.6B), (astrocytes sharing the same code of fluorophores) around the injury site revealed the presence of all type of astrocyte subpopulations: pial, upper- and lower-layers protoplasmic, juxtavascular, and fibrous. Upper- and lower-layers protoplasmic astrocytes were the most prevalent followed by fibrous, while juxtavascular and pial astrocytes represented smaller fractions (Fig. 6C). Quantitative analysis of the number of sibling cells showed the variability in clone size and dispersion across astrocyte subpopulations (Fig.6C). Finally, the analysis of the frequency of astrocytes subpopulations showed that the “strong” type of reactive response (A+B) had the highest proportion of protoplasmic (upper layers) and fibrous astrocytes, while the “very strong” type exhibited the highest proportion of pial astrocytes (Fig.6D). Overall, our data showed a classification to detect morphological similarities and differences among astrocyte subpopulations, suggesting TBI-related morphological signatures in reactive astrocytes.

**Figure 6.**
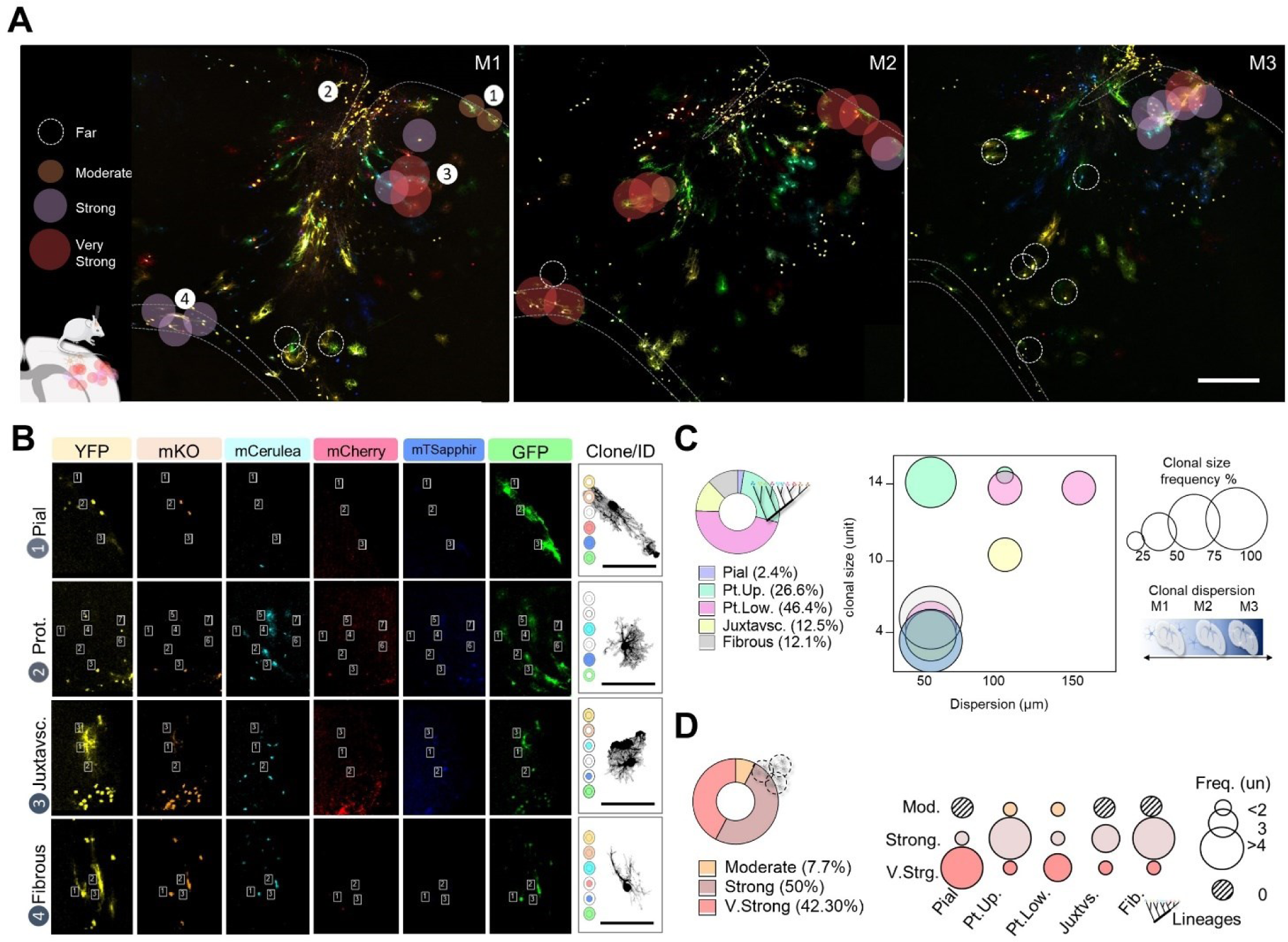
Exploratory analysis of reactive astrocytes responses in animal model of TBI. A. Representative TBI sections, astrocytes distribution and reactive responses. B. Representative images of the clonal analysis in the TBI model. Astrocytes with the same color code composition were assigned as sibling cells (clones). C. Pie chart, and bubble graphs of the frequency and dispersion of sibling cells (clones) among astrocyte subpopulations. D. Bubble graph of astrocytes subpopulation frequency among reactive responses. Scale bar A, 250 μm; and B, 100 μm.

## Discussion

In this work we explored specific subpopulations of StarTrack-labeled astrocytes following traumatic brain injury, with a particular emphasis on their morphology-dependent responses. Our study unveiled the intricate single-cell modifications within critical astrocytic components-soma, branches and branchlets, shedding light on the considerable variability in reactive responses among astrocytes subpopulations. The study of astrocytes encompasses various dimensions including regional, molecular, and biochemical aspects, developmental stages, interspecies adaptations, aging, brain damage, and neurological conditions (Escartin et al., 2019; Khakh et Deneen, 2019; Matias et al., 2019; Pestana et al., 2020; Bayraktar et al., 2020; Falcone et al., 2021; Endo et al., 2022). From pioneering works in glia biology (Virchow 1846; Cajal, 1913; Somjen, 1988) to the most recent state-of-the-art compilation papers (Endo et al., 2022; Baldwin et al., 2023; Verkhratsky et al., 2023), we now understand that the size, shape, and complexity of astrocytes are intimately linked to their functional status and their capacity to interact with diverse neural and non-neural cells.

Here, we identified distinct subpopulations of StarTrack-labeled astrocytes based on functional, regional, and cell-lineage perspectives. According to their distribution across cortical regions and layer-specific interactions, we categorized StarTrack-labeled astrocytes into pial, protoplasmic (both upper- and lower-layers), juxtavascular, and fibrous subpopulations. Studies have demonstrated the critical role of astrocyte morphological variations in their specialized functions. For example, human protoplasmic astrocytes exhibit highly intricate branching morphologies, which are closely linked to the regulation of synaptic communication, while fibrous astrocytes display simpler branching patterns, primarily involved in structural support (Oberheim et al., 2009; Lanjakornsiripan et al., 2018; Verisokin et al., 2021; Degl’Innocenti et Dell’Anno, 2023; Holt., 2023). Moreover, interactions with the vascular network contribute to the presence of perivascular and juxtavascular astrocytes. Perivascular astrocytes, identified as protoplasmic astrocytes extending end-feet to blood vessels, and juxtavascular astrocytes, identified as astrocytes with cell bodies fully associated with blood vessels, are integral to vascular regulation and maintenance (Bardehle et al., 2013; Khakh et Sofroniew, 2015; Götz et al., 2021; Verkhratsky et al., 2021; Ojalvo-Sanz et al., 2024).

Once we established the morphological characteristics of reactive astrocytes, we sought to explore the distinct morphological alterations among astrocytes subpopulations in response to injury. Currently, various measurements are used to assess astrocyte morphological changes that could correlate to their function in response to injury. Unfortunately, most of these analyses encounter limitations due to the inherent constraints of microscopes in resolving parameters such as the overall complexity of 3D branching in astrocytes, including their finer processes like branchlets and leaflets (Baldwin et al., 2023). In this work, and despite encountering interindividual variability values for most of the size-related parameters (CV values), our approach using the StarTrack labeling technique in conjunction with a thoughtful morphological analysis, resulted in a pioneer method of analysis to correlate morphological changes in astrocyte subpopulations such as their shape, size, or a combination of both factors with their function measured as intensity of astrocyte response to an injury. This underscores the significance of considering astrocyte morphology as a multifaceted set of features occurring simultaneously.

Indeed, while shape-related parameters such as “solidity,” “circularity,” and “complexity” are less frequently analyzed compared to size-related parameters, they proved to be highly effective in detecting differential reactive responses among astrocyte subpopulations. These shape-related parameters served as global descriptors, measuring the extent of changes in astrocyte shapes. For instance, in the case of upper and lower protoplasmic reactive astrocytes, we observed lower circularity and solidity ratios, indicating more irregular and spongiform cell bodies, along with enriched Sholl profiles (complexity), which described cell branching density and spatial distribution. These variations likely reflect the adaptation of protoplasmic astrocytes to the characteristic loss of cell polarity and disruption of tissue organization within the damaged neural network at the primary injury site. Following a TBI, many processes such as cell death, neuroinflammation, cell proliferation, and tissue repair overlap to one another, resulting in loss of astrocyte subpopulations and the formation of astrocyte borders, where newly formed astrocytes demarcate and separate damaged areas from the healthy surrounding tissue (Burda et Sofroniew, 2014; Wahane et Sofroniew, 2022; Freire et al., 2023). Interestingly, we found that not only protoplasmic astrocytes, but also pial and juxtavascular astrocytes experienced morphological changes in response to the injury (reactivity) that integrated shape- and size-related parameters. These reactive astrocytes exhibited “more irregular” and “spongiform” cell bodies (lower circularity and solidity values) and increased thickness. The increase in thickness, a size-related parameter, has been associated with the enrichment of cytoskeletal content or cell swelling, typical from the hypertrophy of reactive astrocytes (Schweitzer et Renehan, 1997; Lafrenaye et Simard, 2019). The increase in the cell body volume may be linked to intense modulation of blood-brain and blood-cerebrospinal fluid barriers function (Lafrenaye et Simard, 2019). TBI causes massive rupture of the meninges, blood vessels, and neural tissue, compromising blood-brain barriers and leading to astrocyte reactivity and modulation of vascular permeability and the movement and exchange of molecules. (Iadecola, 2017; Schaeffer et Iadecola, 2021).

Subsequently, we explored the relationships between astrocyte reactivity, and their spatial distribution in upper- and lower-layers protoplasmic astrocytes. Lower-layer protoplasmic astrocytes exhibited a significant reactive response in nine out of the ten parameters evaluated, whereas upper-layer protoplasmic astrocytes displayed a significant change in only one parameter. To delineate areas of reactive gliosis (A1 or contusion area) from GFAP-normal-appearing astrocyte profiles (A2 or pericontusion area), we utilized a GFAP antibody to identify the extension of the lesion in the brain parenchyma. This approach is consistent with previous studies in both human and mouse models, that divided the damaged region to investigate specific cellular and tissue aspects. These investigations have encompassed molecular aspects (Harish et al., 2015; Moeendarbary et al., 2017), vascular and perfusion dynamics (Sword et al., 2013), metabolic changes (Wu et al., 2013), and clinical progression (Adatia et al., 2021).

Our analysis revealed that regardless of the area of study (A1 or A2), upper-layer protoplasmic astrocytes did not exhibit major morphological variations unlike lower-layers protoplasmic astrocytes, which displayed marked, differential and distinctively morphological profiles between both areas. Interestingly, T-SNE and cluster analyses demonstrated that lower-layer reactive astrocytes from A2 and control astrocytes shared similar morphological profiles. At a functional level, since cell morphology is primarily dependent on the reorganization and enrichment of cytoskeletal proteins, the distinct variation in size- and shape-related parameters, particularly in lower-layer protoplasmic astrocytes from A1, suggested a robust response of these subpopulations to the injury in the damaged area. In glial biology, the territorial organization of protoplasmic astrocytes into non-overlapping domains has been well documented, as these domains play crucial roles in regulating neural network function by modulating glia interactions with neural and non-neural cells (Bushong et al., 2002; Ogata et Kosaka, 2002; Wilhelmsson et al., 2006; Pekny et Pekna 2014; Verkhratsky et al., 2021). Previous studies have noted size-related variations in astrocyte domains that are regionally dependent (Wilhelmsson et al., 2006; Verkhratsky et al., 2021), which interact between them by the enrichment of membrane structures at their terminal branches expressing different channels, hemichannels and gap junctions to interchange molecules and signals (Cotrina and Nedergaard, 2012; Lafrenaye et Simard, 2019; Verkhratsky et al., 2021). In this work, we observed that protoplasmic reactive astrocytes at the contusion core (A1) were enriched by primary and secondary branches and leaflets (Sholl intersections profile), that could promote glial physical interactions within astrocyte domains. Our findings indicated that astrocyte distribution among cortical layers, and their proximity to the damaged region (contusion core), influenced their reactive response.

Finally, in this work we are pioneers in establishing a method to classify astrocytes based on their morphology changes in response to injury, correlating different morphological parameters with their intensity of the response. Previous works have introduced this type of approaches to study neuronal subpopulations for example in the nucleus of the solitary tract (Schweitzer et Renehan, 1997) or to categorize diverse subpopulations of hippocampal astrocytes in migratory birds (Carvalho-Paulo et al., 2018; Henrique et al., 2021; de Almeida et al., 2022), reactive microglia in models of encephalitis (de Sousa et al., 2015; da Silva Creão et al., 2021) or in models of neuroinflammation (Fernández-Arjona et al., 2017); and immune cells in starfishes (Karetin, 2021). Here, we developed a multimodality index (MMI) to select parameters that best describe the population heterogenicity, and subsequently applied hierarchical clustering (HC), and principal component analysis (PCA) to identify and construct a dendrogram of reactive responses. Our analysis using a TBI model revealed three first-level categories astrocyte node responses that we named as “moderate,” “strong,” and “very strong” (Fig.5). This approach provided helpful cut-off values for each node level, offering insights for further analyses on the variety of responses of reactive astrocytes. We found an enrichment of sibling protoplasmic astrocytes in both “strong” and “very strong” reactive responses, fibrous, and pial astrocytes, highlighting the importance of lineage-specific responses in brain injury. Our studies demonstrated that sibling astrocytes may establish preferential coupling and form major-like domains (Gutiérrez et al., 2019), which could share electrophysiological properties (Götz et al., 2021). Our analysis sheds some light on the intricate relationship between astrocyte lineages, their clonal origin, and the diverse reactive responses to a traumatic brain injury.

Our morphological reconstructions and categorization of reactive astrocytes provide quantitative insights into the spatial distribution and complexity of reactive astrocytes. In recent years, machine learning tools have demonstrated the value of such datasets for advancing the knowledge of basic science, computational modeling, and translational research (Ascoli et al., 2023). Moreover, while pioneering studies have identified at least two subtypes of reactive astrocytes based on morphological changes (Sofroniew, 2020), here we provide a comprehensive analysis that shows the intricate nature of reactive astrocyte responses and their potential impact on neural network dynamics in the context of brain injury. By elucidating morphology-based reactive responses of astrocyte subpopulations, particularly those at the lesion core, we contribute to a deeper understanding of astrocyte heterogeneity and its implications for neuroinflammation and tissue remodeling in brain injury contexts.

## Methods

### Animals

Animals were handled in compliance with the European Union guidelines on the use and welfare of experimental animals (2010/63/EU), and the Spanish legislation (Ministry of Agriculture, RD 1201/2005 and L32/2007). Procedures were approved by the CSIC Bioethical Committee and the Community of Madrid (Ref. PROEX 274.7.22) in Spain. Cajal Institute (Madrid, Spain). We used isogenic C57Bl/6J mouse pregnant females at the embryonic age 14 (E14) supplied by the stock from the Cajal Institute (Madrid, Spain). Animals were housed in standard cages, maintained under controlled light-dark cycles (12hs-12hs) with food and water *ad libitum*. We made all efforts to minimize suffering and the number of animals used.

### StarTrack mixture and *in utero* electroporation

Cortical astrocytes were labeled using the StarTrack method as previously described (García-Marqués et López-Mascaraque, 2013). StarTrack is a multicolor-based bar code method (cytoplasm and nuclear labeling) that allows to tag single neural progenitors and follow their GFAP+ progeny (glial cells). GFAP StarTrack is based on the PiggyBac system. This system is composed of two elements, one, the DNA transposon vector that carries a gene encoding a fluorescent reporter protein (YFP, mKO, mCerulean, mCherry, Mt-Sapphire, and eGFP) under the control of the human glial fibrillary acidic protein GFAP and the H2B promoter (only for nuclear labeling); and second, the transposase enzyme (CMV-hyPBase) which dimerizes the transposon at the ITR (Inverted Terminal Repeat) sequences, and guide and insert the fragment into areas of transcription (TTAA-AATT) in the genome (Fig,1A). Thus, PiggyBac generates stable gene integration with high rates of transcription. GFAP StarTrack leads to consistent and higher levels of expression of the gene and produces the labeling of the cytoplasm and/or the nucleus of the cells.

Pregnant, C57BL/6j mice (P30-45) in gestational period E14 were deeply anesthetized with an isoflurane vaporizer system, Isova Vet, 2 ml/L, (4% induction, 2-3% maintenance, Centauro, Barcelona, Spain) and after conscious and pain assessment, were dissected the skin and abdominal tissues and exposed the uterine horns and embryos. Each embryo was carefully manipulated to receive an intraventricular injection of 2μl of GFAP Startrack plasmid mixture (2-5mg DNA/ml and 0,1% fast green solution to confirm the site of the injection and diffusion). Next, embryos were electroporated using an Electroporator ECM 830 system (BTX, Massachusetts, US) connected to 5mm tweezer type electrodes (program: 1-2 trains, 5 square pulses, 35V, 50 ms, followed by 950 ms intervals). Finally, uterine horns were placed back, and the abdominal tissues and skin were closed with absorbable suture 4/0 (Surgicryl, Hünningen, Berlin) and silk 4/0 (Lorca-Marin, Murcia, Spain). Postoperative care included antibiotic and analgesic administration (2.27% enrofloxacin, 5 mg/kg, Baytril, Bayer, Kiel, Germany and 0.5mg/ml meloxicam, 300 μg/kg, Metacam, Boehringer Ingelheim Vetmedica GmbH, Rhein, Germany, subcutaneous) in controlled temperature environment (37°C). Thus, neural progenitors of the subventricular zone were labeled with GFAP StarTrack plasmid mixture.

### Model of brain damage by traumatic brain injury

Next, electroporated mice were submitted to a model of brain injury in the somatosensory cortex where GFAP StarTrack astrocytes clones will develop a reactive response (detailed information could be found in Martin-Lopez et al., 2013). For this purpose, adult (P50, n=2 or P90, n=2) GFAP StarTrack mice were deeply anesthetized with an intraperitoneal injection of Equithesin (dose 0.33ml/100g, NIDA Pharmacy, Baltimore, MD, USA) and after conscious and pain assessment, were submitted to unilateral or bilateral penetrating lesion with a 22-gauge (0,7mm) needle in the somatosensory cortex at the following stereotaxic coordinates (relative to Bregma) AP 0 and/or-2mm, ML 3.5 mm, DV - 2mm. Postoperative care included antibiotic and analgesic administration (2.27% enrofloxacin, 5 mg/kg, Bayer, Kiel, Germany and 0.3 mg/ml buprenorphine, 8 µg/kg, Buprex, Merck & Co., Inc., NJ, USA) in a controlled temperature environment (37°C). Finally, three to five days post-lesion (3-5dpl), mice were deeply anesthetized with intraperitoneal injection of sodium pentobarbital (40–50 mg/Kg, Dolethal, Vétoquinol Ltd, Buckingham, UK), transcardially perfused with 4% paraformaldehyde and post-fixed (overnight). Additionally, we used adult (P30, n=2) GFAP StarTrack (mCherry and YFP) mice under physiological conditions as controls. Coronal sections of 50µm were collected.

### Immunohistochemistry

Serial brain sections around the lesion core were rinsed several times with 1xPBS-0.1% Triton (0.1% PBST) and incubated in a blocking solution containing 5% normal goat serum NGS (5% NGS, 0.1% PBST) for 60 minutes, at room temperature. After the time, sections were incubated with selected primary antibodies, rabbit anti-Cux1 (1:300, Proteintech, 11733-1-AP); mouse anti-GFAP (1:500, Millipore/Thermo Fisher Scientific, MAB359); and mouse anti-Nestin (1:100, Cell Signalling, 4760); previously included in blocking solution (5% NGS, 0.1% PBST) overnight, at 2–8°C. The next day, sections were rinsed (0.1% PBST) and incubated with the corresponding secondary antibody, goat anti-rabbit coupled to Alexa Fluor 488 (1:1000, Invitrogen/Thermo Fisher Scientific, A11008); and goat anti-mouse coupled to Alexa Fluor 568 (1:1000, Invitrogen/Thermo Fisher Scientific, A110004) in DAPI solution (1:10000, Sigma Aldrich Corporation, Saint Louis, EUA) for 60-90 minutes, at room temperature. Finally, the sections were rinsed and mounted onto slides with aqueous solution (Fluoromount G, Electron Microscopy Sciences, EUA).

### Image acquisition

Image acquisition was performed at the confocal microscopy unit (Instituto Cajal-CSIC), using Leica TCS-SP5 (Houston, United States) and Leica STELLARIS 8 STED HM (Wetzlar, Germany) confocal microscopes. Imaging of GFAP astroglial populations was performed as described in Martin-Lopez et al., (2013) and Figueres-Oñate et al., (2016). Fluorescent proteins (XFP) were captured in separate channels. The wavelength of excitation (Ex) and emission (Em) for each XFP were: mT-Sapphire, Ex 405nm, Em 520-535 nm, mCerulean, Ex 458 nm, Em 468-480 nm, EGFP, Ex 488 nm, Em 498-510 nm, YFP, Ex 514 nm, Em 525-535 nm, mKO, Ex 514 nm, Em 560-580 nm, and mCherry, Ex 561 nm, Em 601-620 nm (Leica TCS-SP5, 20x). Confocal laser lines were between 25-40% (Leica TCS-SP5, 20x) and the maximum projection images were created using LASAF Leica (v.3.02.16120, Leica Application Suite X, Leica Microsystems, Houston, United States) and ImageJ software (1.49v, http://rsbweb.nih.gov/ij).

### Clonal analysis

Clones of reactive astrocytes around the lesion were defined as those cells sharing fluorescent marks in the nucleus and the cytoplasm. We analyzed 281 cells corresponding to 35 clones of one GFAP StarTrack mouse. Analysis was conducted using ImageJ software (1.49v, http://rsbweb.nih.gov/ij). We analyzed the percentage of clones per astroglial lineage, the quantity (mean), frequency and dispersion of sibling cells per astroglial lineage.

### Morphometric analysis

For morphometric analysis, we analyzed 117 cells of five GFAP StarTrack electroporated mice submitted to brain injury (TBI group) and 45 cells of two GFAP StarTrack (mCherry and YFP) mice in physiological conditions (control group). We used the set measurement options “area” (µm^2^), “convex hull area” (µm^2^), “perimeter” (µm), “circularity” (4π(“area”) ⁄(“perimeter”)^2^, ratio), “solidity” (“area” ⁄ “convex hull area”, ratio). The simple neurite tracer (SNT) tool was used to reconstruct and create a 3D mask of the cell and quantify the following parameters: “number of branches” (units), “length” (µm) “thickness” (relative to 0.05 threshold, µm^3^) “intersections” (units), “radius” (units) or 3D distance” (“4(radius)”, µm), and “complexity” (Sholl intersection profile, “intersections” as a function of “3D distance”, au). Of note, we also included Sholl components “intersections”, “radius” and “complexity”, that correspond with the number of times a branch “intersects” imaginary concentric circles at a given “radius”, centered on the soma (Sholl intersections profile). As a result, Sholl analysis give a one-dimensional profile (“complexity”) of astrocyte branching allowing the comparison of branches density and spatial distribution (Sholl, 1953; Bird et Cuntz., 2019; Baldwin et al., 2023).

### Reactive response categorization

Morphometric parameters were first considered as separated features and next, as being part of reactive astrocytes categories (adapted from Schweitzer et Renehan, 1997). First, multimodal index analysis (MMI) guided the selection of parameters for clustering. MMI formula ([M3^2^+1]/[M4+3{(n-1)^2^/(n-2) (n-3)}]) include skewness (M3) and kurtosis (M4) measures that describe the shape of the data distribution curve. Next, we performed Hierarchical Clustering (HC), Principal Component Analysis (PCA), of the cell population. Values of selected parameters were normalized using the Z-score method, in which the mean of each parameter is subtracted from each value. For HC, Euclidean distance measure calculated the length of the segment connecting two values (2.3 distance threshold). For PCA, we selected principal components (PC) 1 and 2 that collectively explained about 80% of the morphological variance of the datasets. Finally, we created a logical tree of reactive astrocytes response-clusters that included cut-off values corresponding to 95% confidence interval (CI).

### Statistical analysis

In the case of clonal and morphological analysis, we considered absolute (number of sibling cells per astroglial lineage, and 2D and 3D parameters comparisons) and relative values (clones per astroglial lineage, and coefficient of variation). In the case of the T-distributed Stochastic Neighbor Embedding TSNE and clustering (HDBSCAN) analysis we considered perplexity value = 5, and minimum cluster size=10 points. In the case of reactive astrocyte categorization, we considered absolute (clusters comparisons, and 2D and 3D parameters) and relative values (parameters profile, cells, and parameters contribution) for each selected parameter. Depending on the case, the graphs show mean, standard deviation (SD), standard error (SEM), and 95% confidence interval (CI). Statistical significance was evaluated using one-tailed unpaired Student’s t test for 2-group comparison of the means, one-way or two-way ANOVA test for multiple comparisons and pairwise comparisons of means (post-hoc test). Values with a confidence interval of 95% (p < 0.05) were considered statistically significant. Statistical analysis of the data and graphical representations were performed using GraphPad Prism (v 5.0, San

Diego, United States) software, and R packages: Tidyverse (https://cran.r-project.org/web/packages/tidyverse/); Rtsne (https://cran.r-project.org/web//packages/Rtsne/Rtsne.pdf; https://github.com/jkrijthe/Rtsne); DBSCAN (https://cran.r-project.org/web/packages/dbscan/index.html); and Factoextra (https://cran.r-project.org/web/packages/factoextra/index.html).

## Author Contributions

LMDG, ACOS, TKEN, EML - Performed experiments, wrote the original draft, reviewed, and edited the final document and figures. MAP and LLM - Conceptualized the study, advised in the execution of the experiments, and reviewed and edited the final document.

## Acknowledgments

We would like to thank Belen García and Carmen Hernandez from the Imaging and Microscopy Unit, Laudelina Garmendia from the Animal Facility, Cajal Institute for the technical support and Jorge García-Marqués, María Figueres-Oñate, Sonsoles Barriola, Carolina Pernía, and Nieves Salvador members of the Lab, for providing instructive advice and stimulating discussions.

---

**Supplementary Figure 1.**
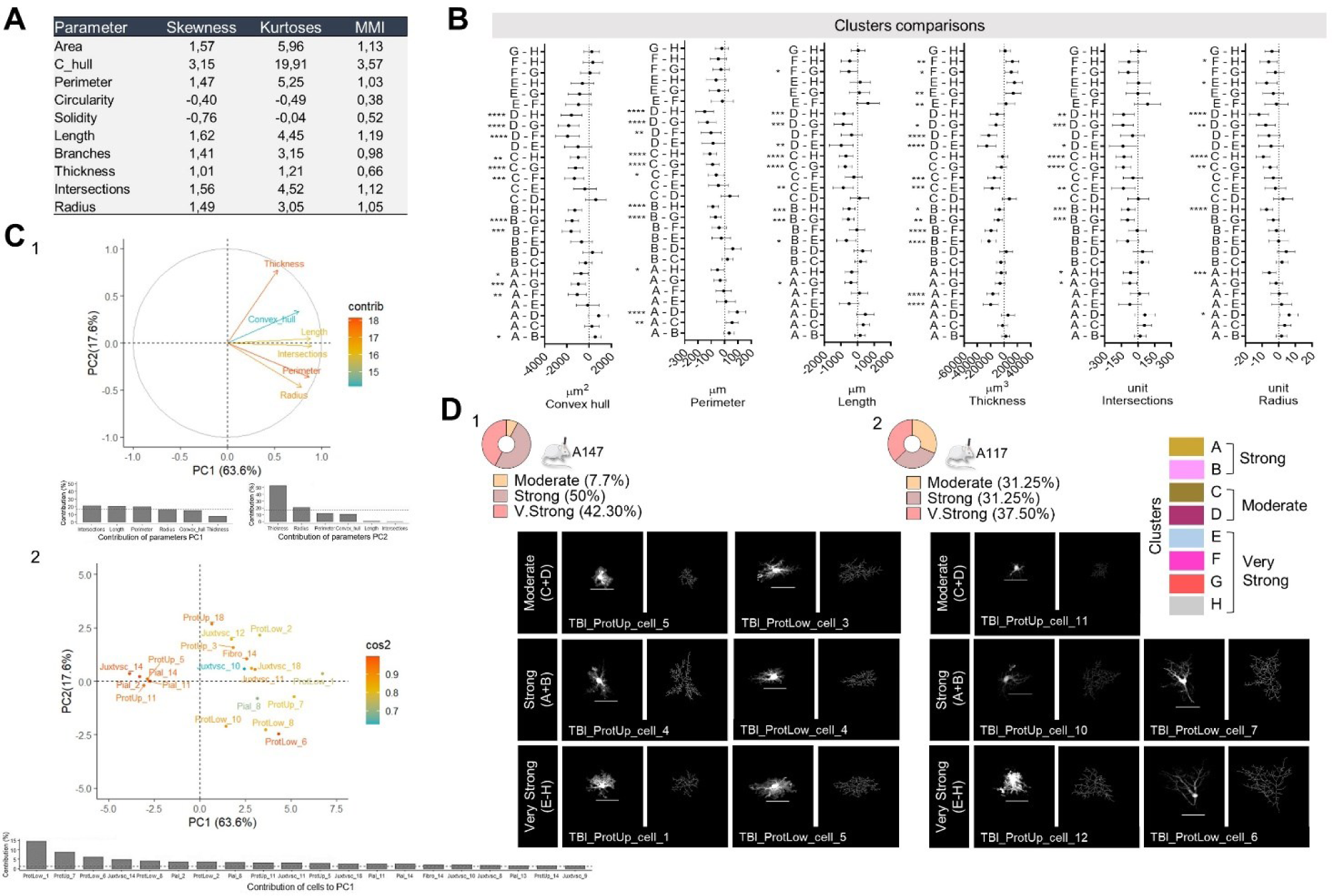
Extended analysis of reactive astrocytes responses. A. 2D/3D parameters multimodal index (MMI). B. Plot of mean differences and standard error (SEM) between clusters in selected parameters. C. Extended PCA plot and graph bar indicating the contribution and importance of selected morphological parameters in the directions of the component values (1); Extended PCA plot and graph bar indicating the contribution and importance of reactive astrocytes (2). Comparison between clusters was performed by ANOVA test, **** p ≤ 0.0001, *** p ≤ 0.001; ** p ≤ 0.01; * p ≤ 0.05. D. Pie charts and representative images of reactive astrocytes responses in two TBI models (A147;1, and A117; 2 mice). Scale bar 50 μm.

