## Supplementary figures and images for "Dissecting Reactive Astrocyte Responses: Lineage Tracing and Morphology-based Clustering"

### Supplementary Figure 1

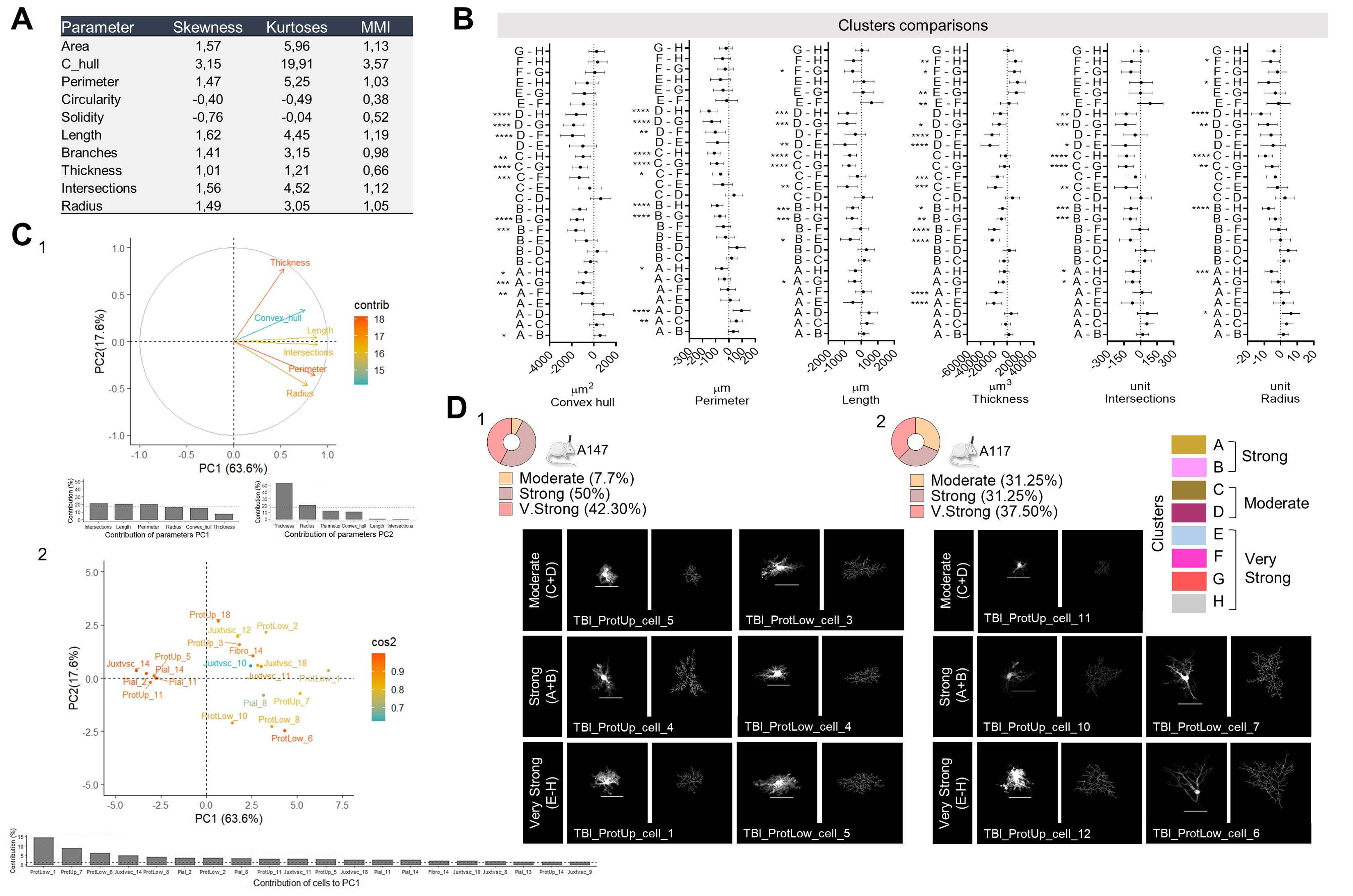
